# Sci-Hi-C: a single-cell Hi-C method for mapping 3D genome organization in large number of single cells

**DOI:** 10.1101/579573

**Authors:** Vijay Ramani, Xinxian Deng, Ruolan Qiu, Choli Lee, Christine M Disteche, William S Noble, Zhijun Duan, Jay Shendure

**Affiliations:** Department of Genome Sciences, University of Washington, Seattle, WA; Department of Pathology, University of Washington, Seattle, WA; Department of Medicine, University of Washington, Seattle, WA; Division of Hematology, University of Washington School of Medicine, Seattle, WA; Institute for Stem Cell and Regenerative Medicine, University of Washington, Seattle, WA; Howard Hughes Medical Institute, Seattle, WA

**Keywords:** Chromatin, Chromosome, Hi-C, Single cell, Single-cell Hi-C, sci-Hi-C, Three-dimensional (3D) genome architecture

## Abstract

The highly dynamic nature of chromosome conformation and three-dimensional (3D) genome organization leads to cell-to-cell variability in chromatin interactions within a cell population, even if the cells of the population appear to be functionally homogeneous. Hence, although Hi-C is a powerful tool for mapping 3D genome organization, this heterogeneity of chromosome higher order structure among individual cells limits the interpretive power of population based bulk Hi-C assays. Moreover, single-cell studies have the potential to enable the identification and characterization of rare cell populations or cell subtypes in a heterogeneous population. However, it may require surveying relatively large numbers of single cells to achieve statistically meaningful observations in single-cell studies. By applying combinatorial cellular indexing to chromosome conformation capture, we developed single-cell combinatorial indexed Hi-C (sci-Hi-C), a high throughput method that enables mapping chromatin interactomes in large number of single cells. We demonstrated the use of sci-Hi-C data to separate cells by karytoypic and cell-cycle state differences and to identify cellular variability in mammalian chromosomal conformation. Here, we provide a detailed description of method design and step-by-step working protocols for sci-Hi-C.

## 1. Introduction

Recent advances in sequencing-based high-throughput genomic technologies, such as Hi-C [1], have revealed that the genome is hierarchically organized in the nucleus of eukaryotic cells [2]. In mammalian cells, this hierarchy of three-dimensional (3D) genome organization is characteristic by multilevel architectural features, such as chromosome territories [3], A/B compartments [1], topologically associated domains (TADs) [4, 5], and long-range chromatin loops between specific loci [6]. The existence of these commonly shared conformational signatures in individual cells has further been demonstrated by recent super-resolution microscopy studies [7, 8]. Moreover, increasing evidence has demonstrated that the 3D genome organization critically impacts various nuclear, cellular and developmental events, such as transcription, DNA replication, cell cycle and cell fate decisions, and that defects in the 3D genome organization can lead to disease phenotypes [9].

However, despite the existence of commonly shared structural features in 3D genome organization, it is important to note that, due to the stochastic nature of the chromatin fiber and the variability in the various nuclear processes (e.g., DNA replication and transcription) between individual cells, the spatial organization of the genome varies considerably, even among cells of a functionally homogeneous population [10]. Hence, the 3D genome organization constructed from ensemble Hi-C represents only an average of millions of cells, but not the actual genome organizations in individual cells. To characterize the 3D genome architecture at the single-cell resolution, we recently developed sci-Hi-C, a massively multiplexed single-cell Hi-C method for mapping the 3D genome in large number of single cells in parallel without requiring physical compartmentalization of each cell or microfluidic manipulation [11]. We have demonstrated that sci-Hi-C is able to separate different cell types on the basis of single cell Hi-C maps and identified previously uncharacterized cell-to-cell heterogeneity in the conformational properties of mammalian chromosomes, enabling “*in silico* sorting” of mixed cell populations by cell-cycle stage[11-13]. We have since improved the initial version of our sci-Hi-C protocol, which is now greatly simplified, easier-to-adapt, and more cost-effective and have successfully applied sci-Hi-C to a diversity of mouse and human cell lines [11-13].

## 2. Method development

Sci-Hi-C [11] was developed by extending the “combinatorial indexing” paradigm, which was first employed in the single-cell ATAC-seq method to measure chromatin accessibility in thousands of single cells per experiment [14]. Similar to ensemble Hi-C protocols, the sci-Hi-C protocol involves cross-linking the cells with formaldehyde. The cells are then permeabilized with their nuclei intact. A 4-cutter restriction enzyme DpnII is used to fragment the chromatin. Nuclei are then distributed to 96 wells, wherein the first barcode is introduced through ligation of barcoded biotinylated double-stranded bridge adaptors. Intact nuclei are then pooled and subjected to proximity ligation, followed by dilution and redistribution to a second 96-well plate. Importantly, this dilution is carried out such that each well in this second plate contains at most 25 nuclei. Following lysis, a second barcode is introduced through ligation of barcoded Y-adaptors. As the number of barcode combinations (96 × 96) exceeds the number of nuclei (96 × 25), the vast majority of single nuclei are tagged by a unique combination of barcodes. All material is once again pooled, and biotinylated junctions are purified with streptavidin beads, restriction digested, and further processed to Illumina sequencing libraries (Fig 1). Sequencing these molecules with relatively long paired-end reads (i.e., 2 × 250 base pairs) allows one to identify not only the genome-derived fragments of conventional Hi-C, but also both rounds of the barcodes, which enable decomposition of the Hi-C data into single-cell contact probability maps. It is worth noting that the throughput (i.e., the number of single cells surveyed per experiment) of sci-Hi-C can readily be scaled up by using 384-well plates to replace the 96-well plates during the two-rounds of barcoding.

**Fig 1.**
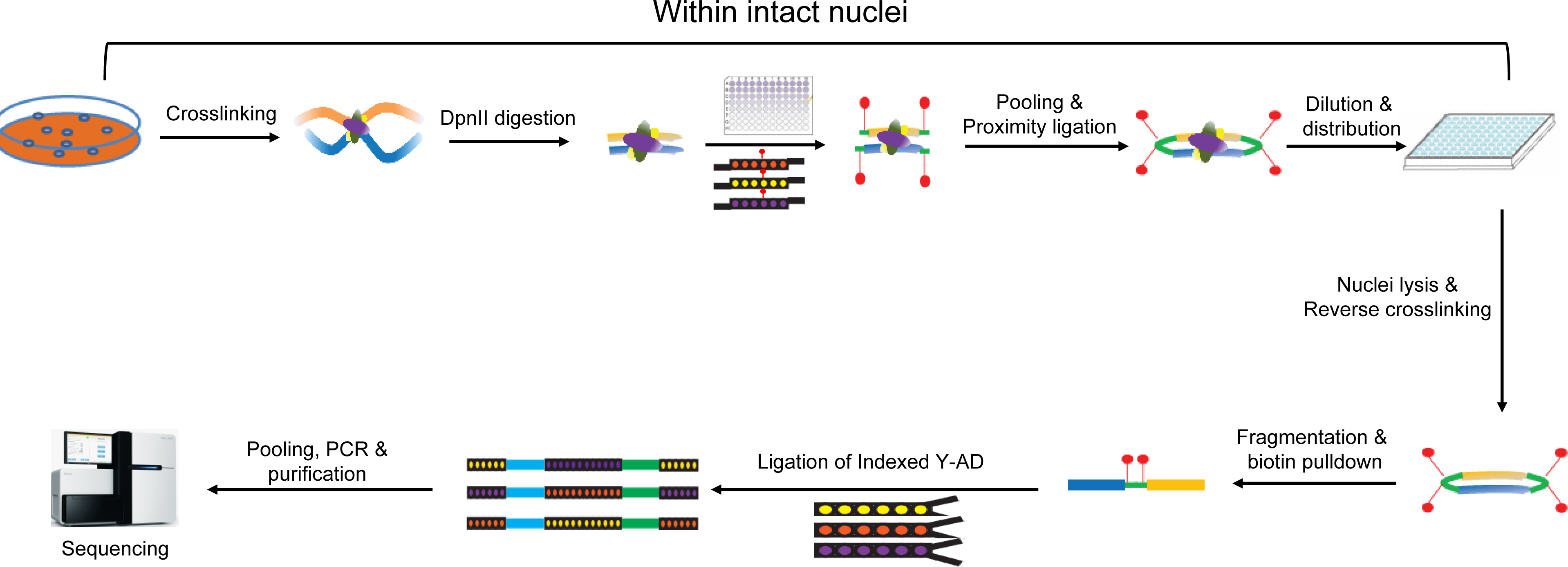
The workflow of sci-Hi-C.

### 2.1. Cell fixation and cell lysis

In this protocol, cells are crosslinked with 2% formaldehyde for 10 min at room temperature (25 °C). However, some cell types (e.g., mouse and human embryonic stem cells) may require different crosslinking conditions and tissues also need to be homogenized. Optimization of crosslinking conditions for a new cell type can be carried out as described previously in [15, 16]. To open the chromatin for enzymatic reactions, while retaining the 3D chromatin conformation and the nuclei intactness, crosslinked cells are then treated sequentially with hypotonic buffer containing IGEPAL CA-630 and buffers containing detergents SDS and Triton X-100. Different cell types may require different SDS concentration, which can be titrated as described previously [15, 16].

### 2.2. Chromatin fragmentation

Sci-Hi-C protocol employs a four-cutter enzyme, DpnII, to fragment the crosslinked chromatin. DpnII digestion is carried out in the presence of detergents (i.e., SDS and Triton X-100) and the size of the majority of the resulting genomic DNA fragments should be less than 1 kb (Fig 2A). It is worth pointing out that not all restriction enzymes are active under these conditions. However, if an enzyme has been used successfully in standard Hi-C protocols, it will likely work for sci-Hi-C. It is also important to note that, since a four-cutter enzyme (e.g., DpnII) has much more cut sites in the genome than a six-cutter enzyme (e.g, HindIII), using a four cutter to fragment the chromatin in principle will generate higher-resolution chromatin interaction maps than using a six cutter.

**Fig 2.**
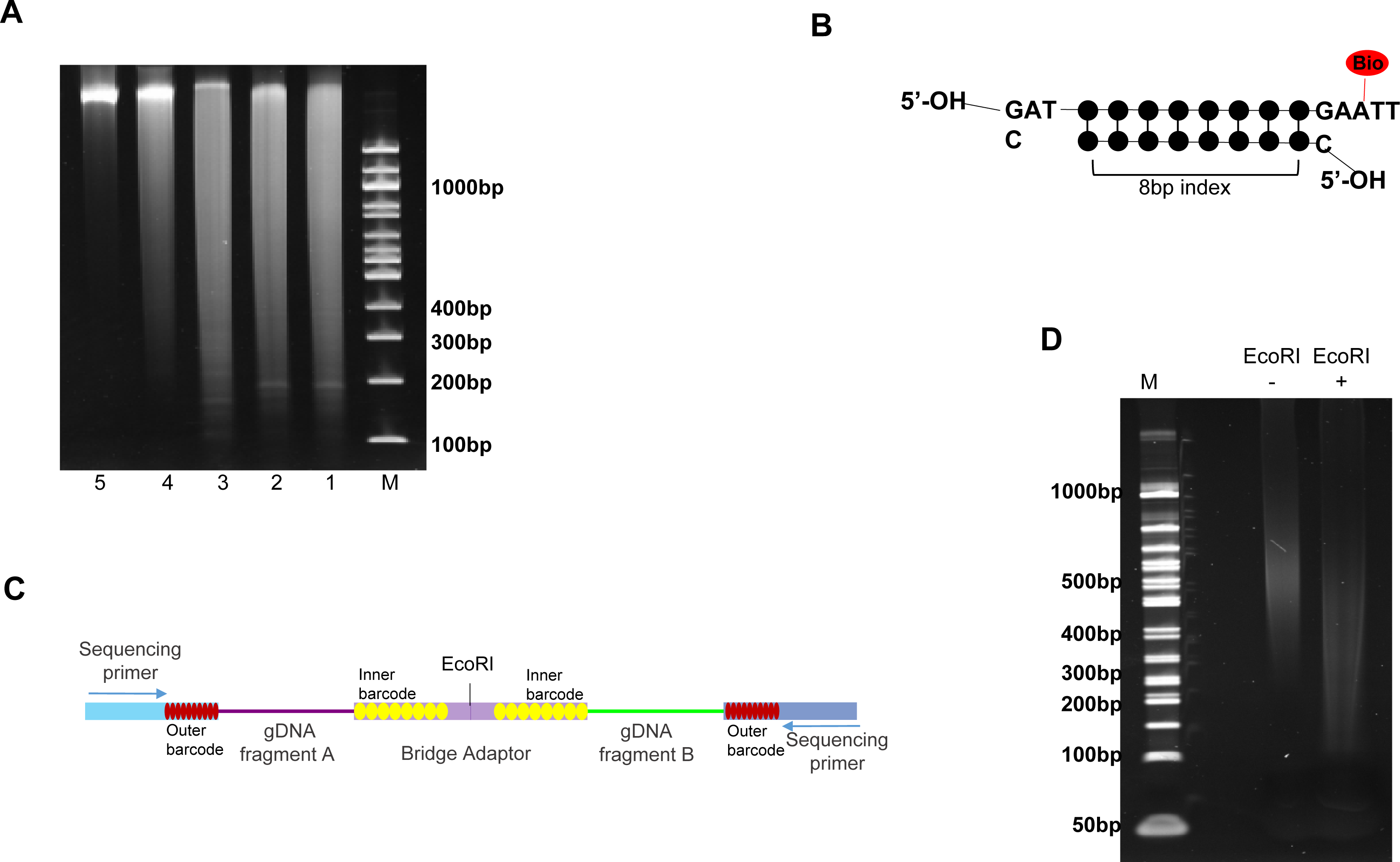
Quality control (QC) of experimental steps in the sci-Hi-C protocol. (A) Example showing the size distribution of genomic DNA samples purified from the nuclei at the indicated various experimental steps of the sci-Hi-C protocol.1-3, genomic DNA isolated from GM12878 (1), HFF (2) and NIH3T3 (3) cells following DpnII digestion (at 37°C for 20 hr), 4, genomic DNA isolated from the nuclei following the ligation of the biotynated- and indexed-bridge adaptors, 5, genomic DNA isolated from the nuclei following the proximity ligation. M, the 100 bp DNA ladder. (B) The structure of the biotinylated- and indexed-bridge adaptors designed for sci-Hi-C. (C) Schematic depiction of the structure of the chimeric DNA molecules in sci-Hi-C libraries. (D) Example of gel electrophoresis showing a sci-Hi-C library before (−) and after (+) EcoRHI digestion. M, the 50 bp DNA ladder. All reactions were run on one 6% TBE-PAGE gel.

### 2.3. Bridge adaptor and 1^*st*^ round barcode design

In the sci-Hi-C protocol, the bridge adaptor is designed to possess several unique features (Fig. 2B). First, it is biotinylated through including a biotin-dT, allowing for enrichment of chromatin interactions with streptavidin-coupled magnetic beads. Second, the two ends of the bridge adaptor are designed to play different roles. One of its ends, through which the adaptor can be ligated to the DpnII digested chromatin fragments, possesses a 5’-GATC overhang. The other end of the adaptor, which is designed to mediate proximity ligation, is not 5′-phosphorylated and its 3’-overhang contains a biotin-dT within a palindromic sequence, which is part of the recognition sequence of EcoRI. Hence, only after the phosphorylation step (Step 37 in the Detailed protocols Section), two adaptor-tagged chromatin fragments can ligate with each other through this end (steps 38-39 in the Detailed protocols Section), which will result in both biotin-labeling and the formation of an EcoRI site) at the junction. And finally, the bridge adaptors harbor the 1^st^ round barcodes, which are drawn from randomly generated 8-mers, such that the following criteria were met: (i) all indexes must have a minimum pairwise Levenshtein distance of 3; (ii) indexes must not contain the sequences TTAA or AAGCTT; (iii) the GC content of all indexes is > 60%; (iv) indexes must not contain homopolymers ⩾ length of 3; and (v) indexes must not be palindromic.

### 2.4. Removal of the excess adaptors after the 1^st^ round barcoding

In the sci-Hi-C protocol, the biotin marking of chromatin fragments and the introduction of the 1^st^ round cellular indexing is achieved simultaneously by ligation of the indexed- and biotinylated-bridge adaptors to the chromatin ends produced by DpnII digestion. It is necessary to clear out the excess free adaptors after the ligation reaction, which otherwise will severely reduce the efficiency of the downstream proximity ligation. We found that multiple rounds of washing of the nuclei with buffers containing SDS and tween-20 can lead to efficient removal of the excess adaptors.

### 2.5. Quality control of sci-Hi-C libraries

During the construction of sci-Hi-C libraries, quality control (QC) analysis should be carried out at several critical steps.

First, the DpnII digestion efficiency can be examined by DNA gel electrophoresis (Fig. 2A). Genomic DNA can be isolated from a small portion (5-10%) of DpnII digested sample, and the size distribution of the DNA fragments determined by running a 1% agarose gel or a precast 6% polyacrylamide DNA gel. Usually an efficient DpnII digestion will result in the majority of DNA fragments with a size<1 kb (Fig. 2A).

Second, the efficiency of the DNA phosphorylation and proximity ligation can also be examined by DNA gel electrophoresis (Fig. 2A). It is worth noting that, the size distribution of the genomic DNAs after proximity ligation should shift significantly towards bigger sizes compared to that of the genomic DNAs isolated from the experimental steps upstream the step of proximity ligation (Fig. 2A).

And finally, as mentioned above, the biotinylated- and indexed-bridge adaptor for marking the chromatin ends in the sci-Hi-C protocol is designed in such a way that an EcoRI enzyme site will be formed at the junction when two chromatin fragments ligate with each other through the bridge adaptors during the proximity ligation. Hence, in sci-Hi-C libraries, each valid DNA fragment is chimeric, consisting of two interacting genomic fragments via the bridge adaptor with an EcoRI site at the joint (Fig. 2C). Therefore, sci-Hi-C libraries can be validated by carrying out an EcoRI digestion assay, i.e., for a valid sci-Hi-C library, there should be an apparent size shift of DNA fragments after EcoRI digestion (Fig. 2D).

### 2.6. Library sequencing

Because the 1^st^ round of indexes are located in the middle of the DNA fragments in a sci-Hi-C library, it requires relatively long paired-end reads to identify these indexes. Therefore, sci-Hi-C libraries are usually sequenced using 250 bp paired-end sequencing on an Illumina Hiseq 2500 instrument.

## 3. Detailed protocols

### 3.1. Adaptor preparation

The sequences of all the adaptors can be found in Ramani et al [11], and all adaptors can be ordered from Integrated DNA Technologies, Inc. (IDT).

1. Setup the 96-well plate for the bridge adaptors (the final concentration is 45 μM),

2. Setup the 96-well plate for the Y adaptors (the final concentration is 20 μM),

| Fill-in mix (add in this order) | Per well |
| --- | --- |
| 10x NEBuffer #2 (NEB, CAT# B7002S) | 10 μl |
| Indexed Forward oligo (100 μM) | 45 μl |
| Indexed Reverse oligo (100 μM) | 45 μl |

3. Anneal the mixtures by heating them to 98 °C for 6 min in a thermocycler, and then turn off the instrument to allow the plates to naturally cool to RT for overnight. Annealed adaptors can be kept at −20 °C.

2. Setup the 96-well plate for the Y adaptors (the final concentration is 20 μM),
| Fill-in mix (add in this order) | Per well |
| --- | --- |
| Water | 50 μl |
| 10x NEBuffer #2 | 10 μl |
| Indexed Forward oligo (100 μM) | 20 μl |
| Indexed Reverse oligo (100 μM) | 20 μl |

### 3.2. Cell lysis and chromatin digestion with DpnII

4. Thaw a vial of 2% formaldehyde-fixed human cells (e.g., HFF c6, GM12878, K562, MIR90, Hela S3 and Hap-1) and mouse cells (e.g., Patski, NIH3T3 and MEFs cells) on 37°C for 10-20 sec.
5. Resuspend in 1 ml of ice-cold cell lysis buffer (10 mM Tris-HCl pH8.0, 10 mM NaCl, 0.2% Igepal CA-630) containing protease inhibitor cocktail (Sigma-Aldrich, Cat#4693132001).
6. Transfer about 2.5 million of each type of cells to a new tube, respectively.
7. Incubate at 0°C for 60 min.
8. Centrifuge for 60 seconds at 850xg at room temperature (RT).
9. Discard the supernatant and resuspend the pellet in each tube in 100 μl SDS buffer (0.5% SDS in 0.1× DpnII buffer (NEB #B7006)).
10. Incubate at 60°C for 10 min.
11. Add 100 μl of water and 25 μl of 10% Triton X-100 to each tube, mix well.
12. Incubate at 37°C for 10 min.
13. Add 40 μl 10× DpnII buffer, 10 μl DpnII (50units/ul, NEB R0543M) and 125 μl water to each tube, mix well.
14. Incubate at 37°C overnight (18-20 hr).
15. Centrifuge for 60 seconds at 850×g at RT.
16. Discard the supernatant and resuspend the pellet in 400 μl water and 3 μl 10mg/ml BSA (ThermoFisher Scientific, AM2616) in each tube, mix well.
17. Centrifuge for 60 seconds at 850×g at RT.
18. Discard the supernatant and resuspend the pellet in 600 μl water and 6 μl 10mg/ml BSA in each tube, mix well. (Note, the volume for cell resuspension dependent on the cell number and the number of 96-well plates used in the next step; the number listed here is for 2 96-well plates).

### 3.3. Ligation of the biotin-labeled, barcoded Bridge adaptors

(∼25,000-100,000 cells /per well, 10 μl reaction volume, 2-4 96-well plates)

19. Distribute T4 DNA ligase/buffer to each well of 2-4 plates (below is for 2 plates, 3 μl per well),

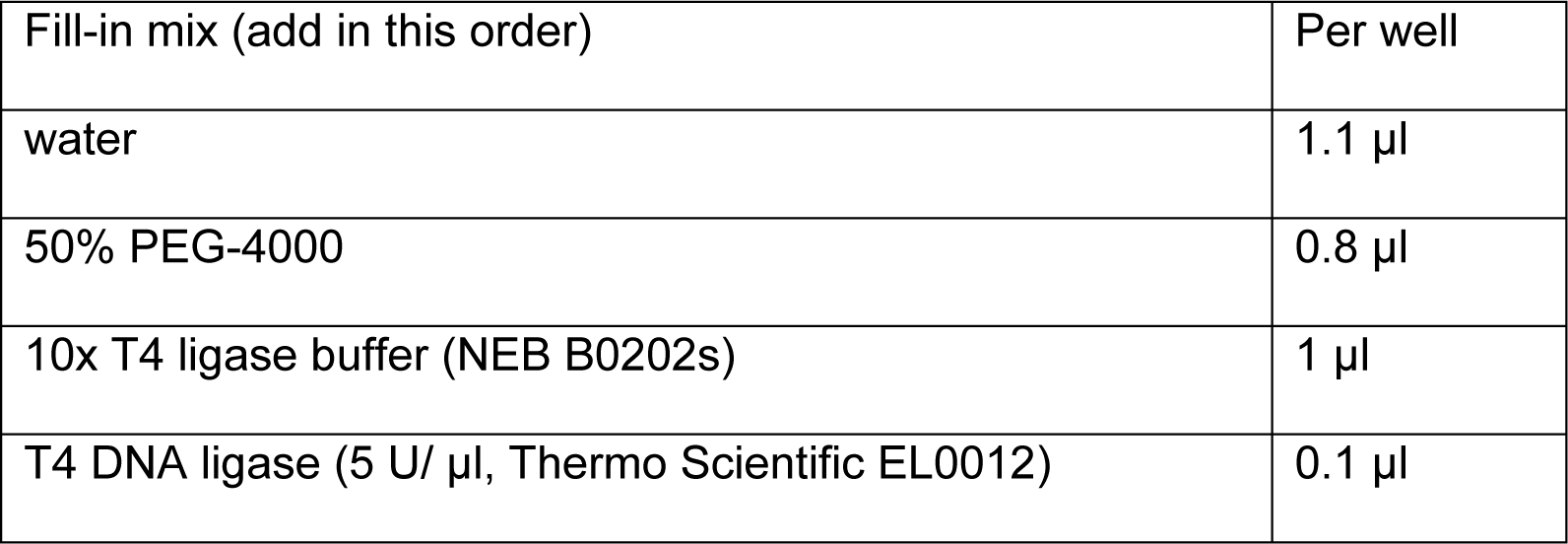
20. Distribute the Bridge adaptor to each well of a 96-well plate using a multi-channel pipette, 2 μl per well.
21. Distribute 5 μl of the above DpnII digested nuclei to each well as below: human cells in one 96-well plate and mouse cells in the other plate. If there are more than two cell types in each specie, the cells can be distributed into different rows of the 96-well plate.
22. Incubate at 16°C overnight.
23. Distribute 1 μl 0.25 M EDTA to each well to stop the reaction.
24. On ice, pool the samples from each well to two 1.5 ml tubes.
25. Centrifuge at 850×*g* at RT for 90 seconds.
26. Resuspend the pellet in 294 μl of water, 3 μl 10%SDS, and 3 μl 10mg/ml BSA, combine the tubes into one.
27. Incubate at RT for 2 min.
28. Centrifuge at 850×*g* at RT for 60 seconds.
29. Resuspend the pellet in 295.5 μl of 2 mM MgCl_2_, 1.5 μl 2% tween-20, and 3 μl 10mg/ml BSA.
30. Centrifuge at 850×*g* at RT for 60 seconds.
31. Repeat steps 29 and 30 one more time.
32. Resuspend the pellet in 294 μl of 2 mM MgCl_2_, 3 μl 10%SDS, and 3 μl 10mg/ml BSA.
33. Centrifuge at 850×*g* at RT for 60 seconds.
34. Repeat steps 32 and 33 one more time.
35. Resuspend the pellet in 294 μl of 2 mM MgCl_2,_ 3 μl 10%SDS and 3 μl 10mg/ml BSA, mix well.

### 3.4. Phosphorylation and proximity ligation

36. Prepare the PNK-ligation mastermix as follows:

37. Incubate at 37°C for 1 hr. Three reactions may be carried out in parallel.
38. Add the following to the above tube,

|  |  |
| --- | --- |
| Fill-in mix (add in this order) | Per reaction |
| water | 50 μl |
| Nuclei | 20 μl |
| 10x T4 ligase buffer | 10 μl |
| PNK (10 U/ μl, Thermo Scientific, EK0032) | 15 μl |
| 10% Triton X-100 | 5 μl |

39. Mix well, incubate at RT for 4 hr.
40. Add 5 μl 10% SDS to each reaction.

|  |  |
| --- | --- |
| water | 172 μl |
| 10x T4 ligase buffer | 20 μl |
| T4 DNA ligase (5 U/ μl) | 5 μl |
| 10 mg/ml BSA | 3 μl |

### 3.5. Serial dilution and reverse cross-linking

41. Carry out serial dilution as follows: transfer 100 μl solution from step 40 to 885 μl water plus 10 μl 10 mg/ml BSA and 5 μl10% SDS, mix well. Transfer 100 μl of the diluted nuclei to 885 μl water plus 10 μl 10 mg/ml BSA and 5 μl10% SDS, mix well. Then transfer 200 μl of the diluted nuclei to 196 μl water plus 100 μl 10x NEBuffer 2 and 4 μl 10 mg/ml BSA, mix well. Now there are about 2-4 cells per 1 μl solution.
42. Distribute 5 μl per well to a low-DNA binding 96-well PCR plate (Bio-Rad, HSS-9601).
43. Incubate at 65°C for overnight. Plates can then be stored at −20°C for future use.

### 3.6. Genomic DNA digestion with AluI + HaeIII

44. add the following to each well of the 96-well plate from the step 43:

45. Briefly spin the plate at 1000 rpm.
46. Incubate at 37 °C for 24 hrs.
47. Incubate at 65°C for 30 min

| Fill-in mix (add in this order) | Per well |
| --- | --- |
| Water | 4.3 μl |
| 10% Triton X-100 | 0.1 μl |
| *AluI* (10 unit/ μl, NEB R0137L) | 0.3 μl |
| *HaeIII* (10 unit/ μl, NEB R0108L) | 0.3 μl |

### 3.7 dA-tailing

48. Add the following dA-tailing reaction reagent to each well of the 96-well plate:

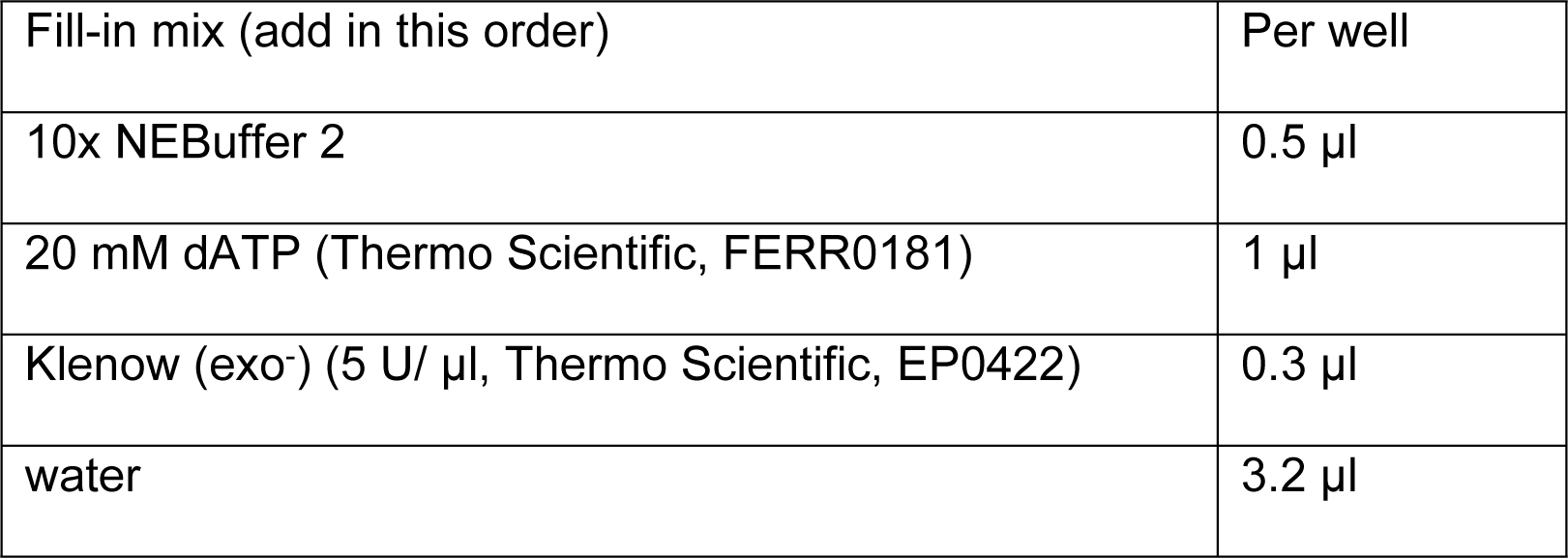
49. Briefly spin the plate at 1000 rpm.
50. Incubate at 37 °C for 45 min.
51. Incubate at 75°C for 20 min.

### 3.8. Ligation of the indexed Illumina Y adaptors

52. Distribute 1 μl Indexed Y adaptor (20 μM) to each well of the 96-well plate.
53. Add the following ligation reaction reagent (14 μl) to each well of the 96-well plate, mix well:

54. Briefly spin the plate at 1000 rpm.
55. Incubate at 16°C overnight.
56. Incubate at 70°C for 10 min

| Fill-in mix (add in this order) | Per well |
| --- | --- |
| water | 7.8 µl |
| 10x T4 ligation Buffer | 3 µl |
| 50% PEG-4000 | 3 µl |
| T4 DNA ligase (5 U/ µl) | 0.2 µl |

### 3.9. Ampure beads Purification and Biotin pull down

57. For each 96-well plate from Step 56, dilute 2.5 ml Ampure beads (Beckman Coulter, A63881) with 3.5 ml 20%PEG-8000 buffer (20% PEG-8000 (Sigma-Aldrich, 89510-250G-F) in 2.5M NaCl), Mix well.
58. Add 50 μl of the diluted Ampure beads to each well of the 96-well plate from the step 56.
59. Combine every 8 wells (each column) into a 1.5 ml low-binding tube, and 12 tubes for a 96-well plate.
60. Rinse each 8-wells with 60 μl EB. Now, the volume in each tube is 700 μl.
61. Incubate at RT for 2 min.
62. Wash two times with 80% Ethanol, combine the 12 tubes into one tube.
63. Elute DNA from the beads with 100 μl EB (10 mM Tris-HCl pH 8.5).
64. Bind to 10 μl DynaBeads M-280 (Thermo Fisher Scientific, 11205D) in 200 μl 1× B&W buffer (5 mM Tris-HCl pH8.0, 0.5 mM EDTA, 1 M NaCl).
65. Incubate at RT for 20 min.
66. Wash 4 times with 200 μl 1× B&W buffer containing 0.1% tween-20.
67. Wash twice with 200 μl EB.
68. Resuspend the beads in 40 μl EB in each tube.

### 3.10. Library amplification and purification

69. Set up 10 PCR reactions as below,

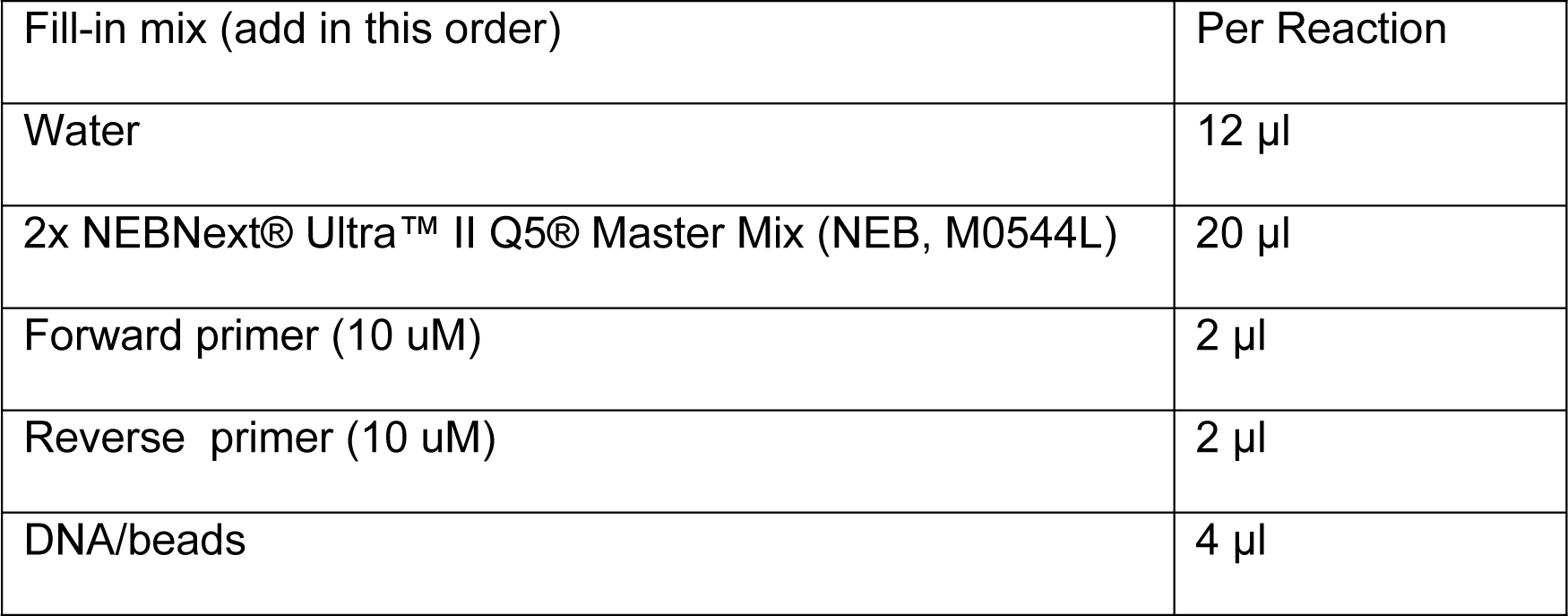
70. Carry out the following PCR program:
71. Using the following PCR program:

72. Pool all the samples from each PCR reaction into a 1.5 ml tube.
73. Add 100 μl Ampure beads and 300 μl of 20% PEG-8000 buffer to get the ratio of 1:1, mix well.
74. Incubate for 5 min at RT.
75. Wash once with 1 ml 80% Ethonal.
76. Resuspend the beads in 100 μl water and transfer the DNA to a new tube.
77. Add 80 μl 20 % PEG solution, mix well.
78. Incubate for 5 min at RT.
79. Wash twice with 0.4 ml 80% ethonal.
80. Repeat steps 76-79 one more time if necessary.
81. Elute DNA with 30 μl water or EB.
82. Determine the DNA concentration using Qubit dsDNA HS kit (ThermoFisher Scientific, CAT# Q32851) according to manufacturer’s instructions.

70. Carry out the following PCR program: 71. Using the following PCR program:
| Cycle number | Denature | Anneal | Extend |
| --- | --- | --- | --- |
| 1 | 98°C, 2 min |  |  |
| 4 | 98°C, 20 sec | 60°C, 20 sec | 72°C, 1 min |
| 8-12 | 98°C, 20 sec | 65°C, 20 sec | 72°C, 1 min |

### 3.11. Library validation by EcoRI digestion

83. Digest a small aliquot of the final sci-Hi-C library with EcoRI to estimate the portion of molecules with valid biotinylated junctions.

84. Incubate at 37 °C for 20 min.
85. Run the entire volume of the reaction on a 2% agarose gel (Fig. 3C).

|  |  |  |
| --- | --- | --- |
| Reagent | Experimental | Control |
| 10x Fast digestion buffer | 1 µl | 1 µl |
| Hi-C product | 1-2 µl (5-10 | 1-2 µl (5-10 ng) |
| Fast digestion EcoRI (ThermoFisher Scientific, FD0274) | 1 µl | 0 µl |
| Water | to 10 µl | to 10 µl |

**Fig 3.**
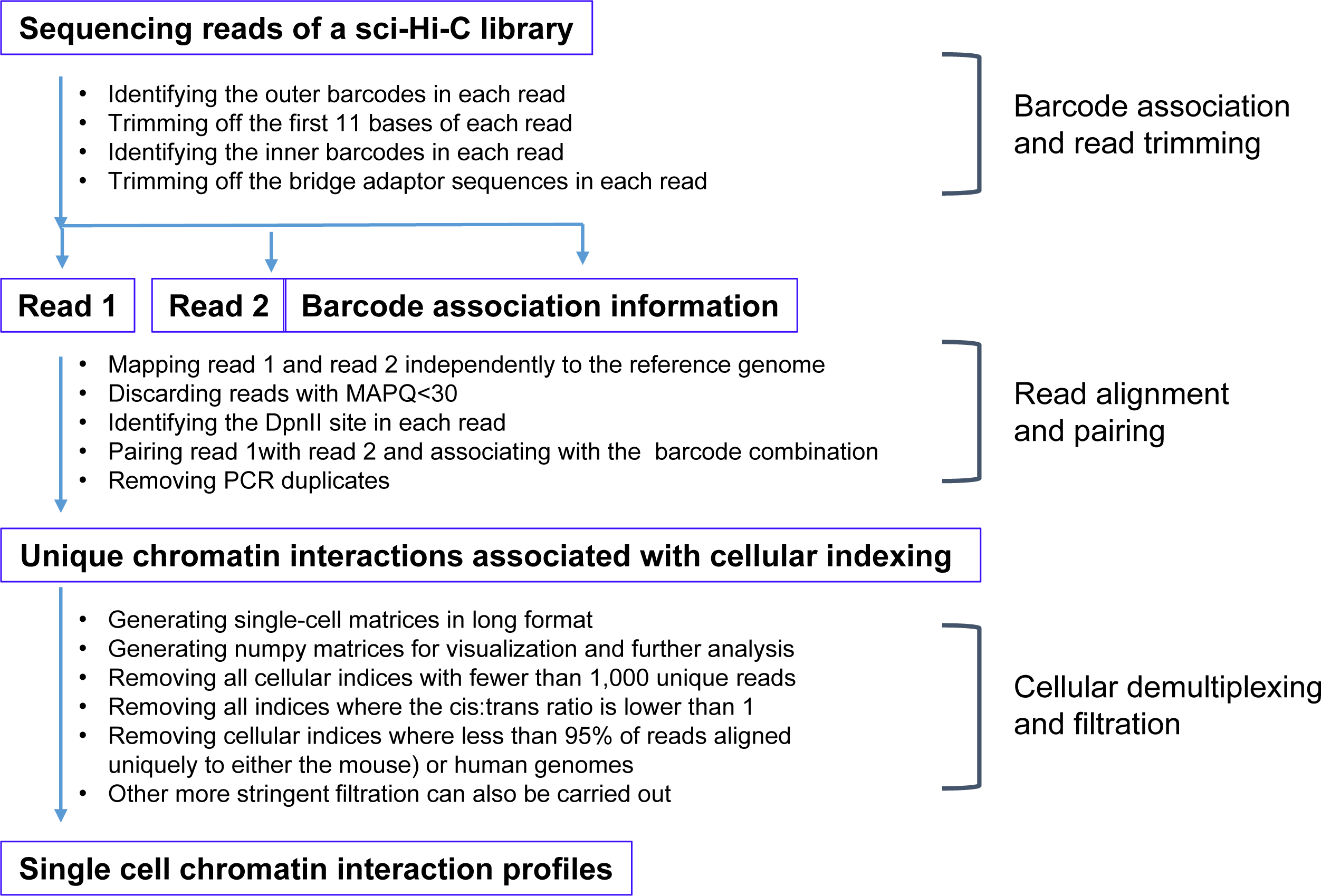
The outline of sci-Hi-C data processing.

## 4. Data processing and expected results

### 4.1. Sci-Hi-C processing pipeline

We developed a computational pipeline to process sci-Hi-C data, which is available at https://github.com/VRam142/combinatorialHiC. This pipeline consists of scripts for conducting the following four types of tasks: (i) barcode identification and read trimming; (ii) read alignment, read pairing, barcode association and filtering of spurious reads; (iii) cellular demultiplexing and quality analysis; and (iv) filtration of cellular indices. The rationale behind these scripts and step-by-step instructions are described in detail in the online methods of Ramani et al [11]. The processing workflow is also outlined in Fig 3.

The quality of a sci-Hi-C dataset can be gauged in several ways. First, the overall distribution of chromatin interactions in a sci-Hi-C dataset should resemble that in a conventional bulk Hi-C dataset (e.g., showing strong intrachromosomal interaction enrichment) (Fig 4A), and thereby the aggregated chromatin contact heatmap created from assembled single cells should be highly similar to that generated from bulk Hi-C assays with the same cell population [11]. Second, in a successful sci-Hi-C dataset, at least forty percent of the read pairs are associated with appropriate barcoding combinations (i.e., cellular indices). Third, if the nuclei remain intact through the experimental steps 4-39 during a sci-Hi-C assay, which can be examined by phase-contrast microscopy, there should be minimal crosstalk between cellular indices (Fig 4B). And fourth, each 96-well plate in a sci-Hi-C experiment can typically yield hundreds to 2000 single cells with at least 1000 unique long-range chromatin interactions (ranging from 1000 to near 1 million interactions).

**Fig 4.**
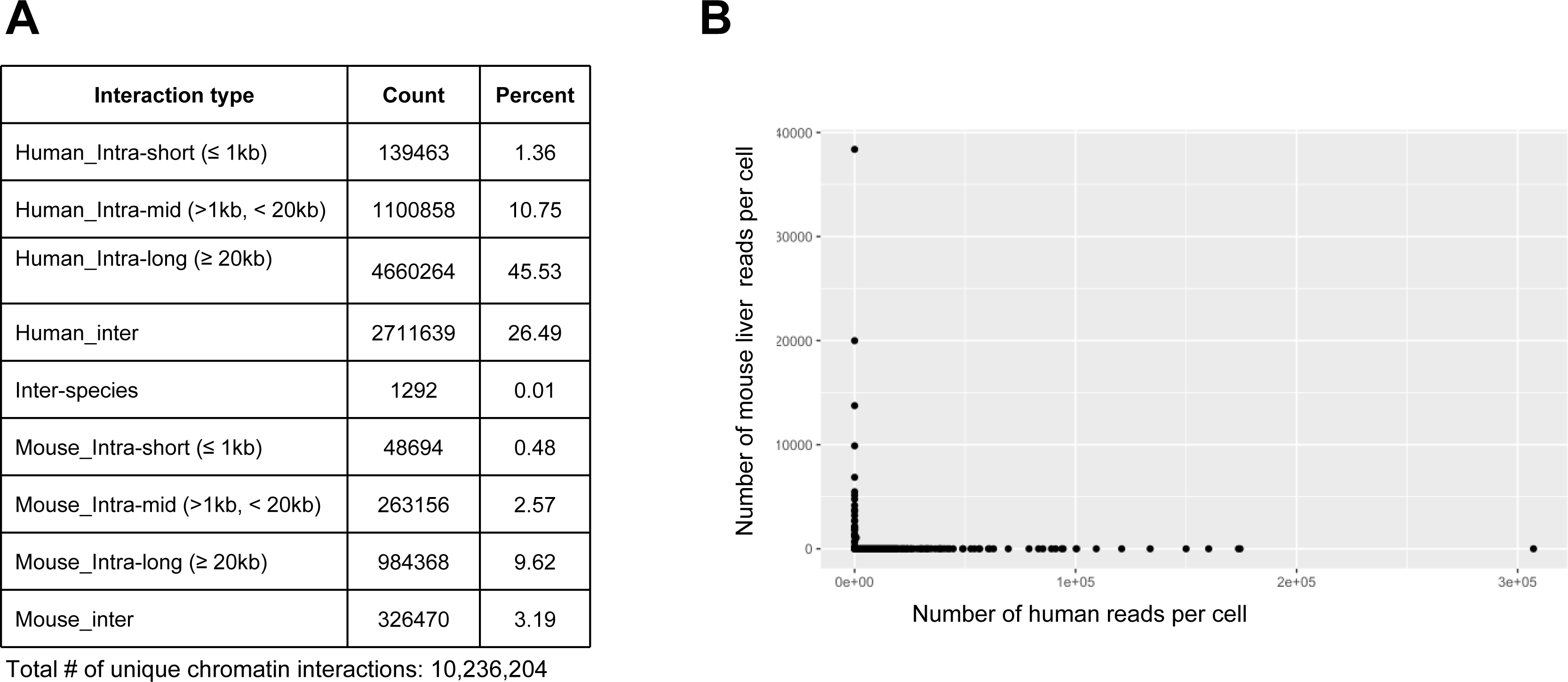
Examples of quality analysis of sci-Hi-C datasets. (A) Example of the chromatin interaction distribution of a sci-Hi-C library containing human and mouse cells. (B) Example scatterplot showing number of reads mapping uniquely to human or mouse genome for individual barcode combinations, indicating low levels of collision between species.

### 4.2. Using sci-Hi-C datasets to characterize 3D genome organization in individual cells

Like other single-cell Hi-C methods [17-20], sci-Hi-C captures snapshots of chromatin conformations from individual cells at the time of cell fixation. Despite the sparsity in genome coverage, single-cell Hi-C assays enable to interrogate 3D genome organization in individual cells. Compared to other single-cell Hi-C datasets, sciHi-C datasets are relatively more sparse yet rich in cell numbers, allowing to obtain single-cell and bulk measurement at once (the latter generated by summing large number of single cells).

We have demonstrated that sci-Hi-C is able to identify cell-to-cell heterogeneity in the conformational properties of mammalian chromosomes and to separate different cell types on the basis of karytoypic and cell-cycle state differences, enabling “*in silico* sorting” of mixed cell populations by cell-cycle stage [11]. Moreover, by combining HiCRep [21] with multidimensional scaling (MDS), we was able to implement an analytical tool to embed sci-Hi-C data into a low-dimensional space, which successfully separate cell subtypes from a cell population based on cell-to-cell variations in cell-cycle phase [13]. More recently, We found that topic modeling [22] provides a powerful tool that is able to capture cell type-specific differences in chromatin compartment structures from sci-Hi-C data [12].

## 5. Conclusions

Sci-Hi-C enables profiling of chromosome conformation in large number of single cells by employing combinatorial cellular indexing.

## Acknowledgements

This work was funded by grants from the NIH (5T32HG000035 to VR, DP1HG007811to JS and U54DK107979 to XD, CMD, WSN, ZD and JS), and ASH Bridge Fund and UW Bridge Fund to ZD. JS is an Investigator of the Howard Hughes Medical Institute.

## High lights

- Sci-Hi-C is based on the concept of combinatorial indexing
- Sci-Hi-C is a high throughput single-cell Hi-C method for mapping chromatin interactiomes in large number of single cells
- Sci-Hi-C does not require special equipment to physically isolate individual cells
- Sci-Hi-C enables “in silico” cell sorting based on cell-to-cell heterogeneity in the conformational properties of mammalian chromosomes

